# An extracellular redox signal triggers calcium release and impacts the asexual development of *Toxoplasma gondii*

**DOI:** 10.1101/2021.02.04.429728

**Authors:** Eduardo Alves, Henry J. Benns, Lilian Magnus, Caia Dominicus, Tamás Dobai, Joshua Blight, Ceire J. Wincott, Matthew A. Child

## Abstract

The ability of an organism to sense and respond to environmental redox fluctuations relies on a signaling network that is incompletely understood in apicomplexan parasites such as *Toxoplasma gondii*. The impact of changes in redox upon the development of this intracellular parasite is not known. Here, we provide a revised collection of 58 genes containing domains related to canonical antioxidant function, with their encoded proteins widely dispersed throughout different cellular compartments. We demonstrate that addition of exogenous H_2_O_2_ to human fibroblasts infected with *T. gondii* triggers a Ca^2+^ flux in the cytosol of intracellular parasites that can induce egress. In line with existing models, egress triggered by exogenous H_2_O_2_ is reliant upon both Calcium-Dependent Protein Kinase 3 and diacylglycerol kinases. Finally, we show that the overexpression a glutaredoxin-roGFP2 redox sensor fusion protein in the parasitophorous vacuole severely impacts parasite replication. These data highlight the rich redox network that exists in *T. gondii*, evidencing a link between extracellular redox and intracellular Ca^2+^ signaling that can culminate in parasite egress. Our findings also indicate that the redox potential of the intracellular environment contributes to normal parasite growth. Combined, our findings highlight the important role of redox as an unexplored regulator of parasite biology.

## Introduction

*Toxoplasma gondii* is a single-cell obligate intracellular parasite from the Apicomplexa phylum that can infect any warm-blooded animal. Its seroprevalence is estimated at more than one-third of the human population^(1, 2)^. Within the host, the asexual lifecycle of *T. gondii* exists as two distinct stages: rapidly proliferating tachyzoites that characterize the acute infection, and slower replicating encysted bradyzoites that are associated with chronic infection^(3, 4)^. While infections are usually benign, in immunocompromised patients and foetuses^(5, 6)^ the lytic tachyzoite lifecycle is responsible for severe clinical pathology. During a lytic cycle, tachyzoites attach onto and actively penetrate host cells, forming a permissive replication niche called the parasitophorous vacuole (PV). Parasites then replicate by endodyogeny^(7)^ until they eventually egress from the host cell, leading to its lytic destruction. The interconversion of virulent tachyzoites with persistent encysted bradyzoites is influenced by host immune pressure and is key to understanding disease recrudescence^(8)^. The ability of *T. gondii* to infect diverse host species relates to its remarkable capacity to resist host defences, efficiently recognize and quickly respond to myriad environmental clues. Among biological signal cascades activated by external clues, Ca^2+^ signaling is the most notable and best studied in *T. gondii* (reviewed in ^(9)^). Ca^2+^ is a ubiquitous signaling molecule with a vital role in tachyzoite host-cell invasion^(10, 11)^, motility^(12)^ and egress^(13)^ through activation of effector proteins such as the plant like Ca^2+^-dependent Protein Kinase 1 (CDPK1)^(14, 15)^ and CDPK3^(16)^. Alongside Ca^2+^, other intracellular signaling molecules are known to play important roles in the tachyzoite lytic cycle. These include phosphatidic acid (PA)^(17)^, and the activation of protein kinase G by cyclic guanidine monophosphate (cGMP)^(18)^.

Contrasting with Ca^2+^, the study of reactive oxygen species (ROS) as signaling molecules is incipient in the Apicomplexa. Hydrogen peroxide (H_2_O_2_) is a neutral ROS molecule with an ability to cross membranes^(19)^, and its roles in signaling are diverse. These include its interaction with glutathione (GSH), and the oxidation of cysteine residues leading to allosteric changes in a variety of proteins such as phosphatases, transcription factors and ion channels^(20)^. Typically, the oxidation of redox-sensitive cysteines is a reversible process catalysed by enzymes that use GSH or nicotinamide adenine dinucleotide phosphatase (NADPH)^(21-24)^ as redox cofactors. Most studies of H_2_O_2_ and *T. gondii* tachyzoites have focused on the ability of host cells to use the damaging oxidative properties of ROS as a component of innate defence, and strategies employed by tachyzoites to overcome this defence^(25-29)^.

A *T. gondii* orthologue of the H_2_O_2_-detoxifying enzyme catalase is expressed in the parasite cytosol, and confers protection against host oxidative stress^(26, 30)^. In 2004 Ding *et al*., ^(26)^ compiled a group of 14 genes related to the *T. gondii* antioxidant system. These typically localized to the parasite cytosol and mitochondria, and were assigned to one of five major redox system classifications: (1) metabolic genes (e.g. superoxide-dismutase and catalase); (2) thioredoxins (Trxs, proteins that promote cysteine thiol-disulfide exchange); (3) Protein Disulfide Isomerases (PDIs, proteins that disrupt or form cysteine disulfide bonds to assist protein folding); (4) glutaredoxin-glutathione (Grx-GSH, small proteins that use GSH as a cofactor for thiol-disulfide exchange) and (5) peroxiredoxins (Prxs, enzymes that detoxify hydroperoxides like H_2_O_2_ and organic hydroperoxides).

Expanding this broad description of the parasite’s antioxidant system, other studies have tested the association of redox with *T. gondii* signaling and cell cycle regulation. Exogenous treatment of intracellular parasites with the reducing agent dithiothreitol (DTT) triggers Ca^2+^ mobilization and parasite egress^(31)^. This egress response was triggered by the depletion of host-cell ATP resulting from the activity of an exported parasite ATPase^(32)^. In a separate study, the oxidation-sensitive protein *Tg*DJ-1 was found to associate with CDPK1 and promote microneme secretion in *T. gondii*^(33)^. More recently, oxidative stress (generated by sodium arsenite) has been shown to trigger tachyzoite differentiation into bradyzoites following phosphorylation of *T. gondii* eIF2α (TgIF2α) by the translation initiation factor kinase *Tg*IF2K-B^(34)^. Together, these data suggest that *T. gondii* modulates biological processes in response to changes in redox homeostasis.

Here, we use an *in silico* approach to establish a compendium of redox-associated genes and provide an updated view of these genes in *T. gondii*. We then investigate the impact of H_2_O_2_ upon parasite biology, demonstrating that a H_2_O_2_ signal outside the boundaries of the infected host cell can be received and interpreted deep within the cytosol of intracellular parasites. We find that exogenous treatment of H_2_O_2_ triggers mobilization of Ca^2+^ culminating in CDPK3-dependent egress. Finally, we use a genetically encoded redox reporter to dissect redox oscillations prior to egress. Unexpectedly, we discover that overexpression of the active catalytic domain of a human Grx in the parasite’s cytosol or PV delays parasite asexual replication. Our results corroborate the existence of a rich variety of antioxidant proteins located in multiples cellular compartments and highlight importance of redox in the basic biology of *T. gondii*.

## Results

### An updated compendium of T. gondii redox-associated genes

We initially sought to update our understanding of the antioxidant response and redox-signaling network in *T. gondii*. We mined gene sequence and annotation information, as well as proteomic datasets present on ToxoDB^(35)^ to update the list of 14 redox associated genes previously summarised by Ding *et al*., 2004^(26)^. Our bioinformatic approach screened for genes containing at least one functional domain of the major redox signaling groups (Trxs, PDIs, Grx-GSHs and Prxs). We identified a total of 58 redox-associated genes (Supplemental material table S1), including 26 Trxs, 16 GRX-GSHs, six PDIs, six Prxs, and four metabolic genes (including three distinct superoxide dismutases and one catalase). With the exception of the metabolic genes^(26, 30, 36, 37)^, the majority of genes representing the other redox signaling groups remain uncharacterized. For an improved view of the subcellular distribution of these gene products, we extracted their primary localisation from the recently published hyperLOPIT dataset^(38)^ (Figure 1 – table supplement 1). This indicated a broad distribution of these proteins throughout the cell and suggested the existence of spatially distinct mechanisms of redox regulation for different subcellular compartments.

**Figure 1:**
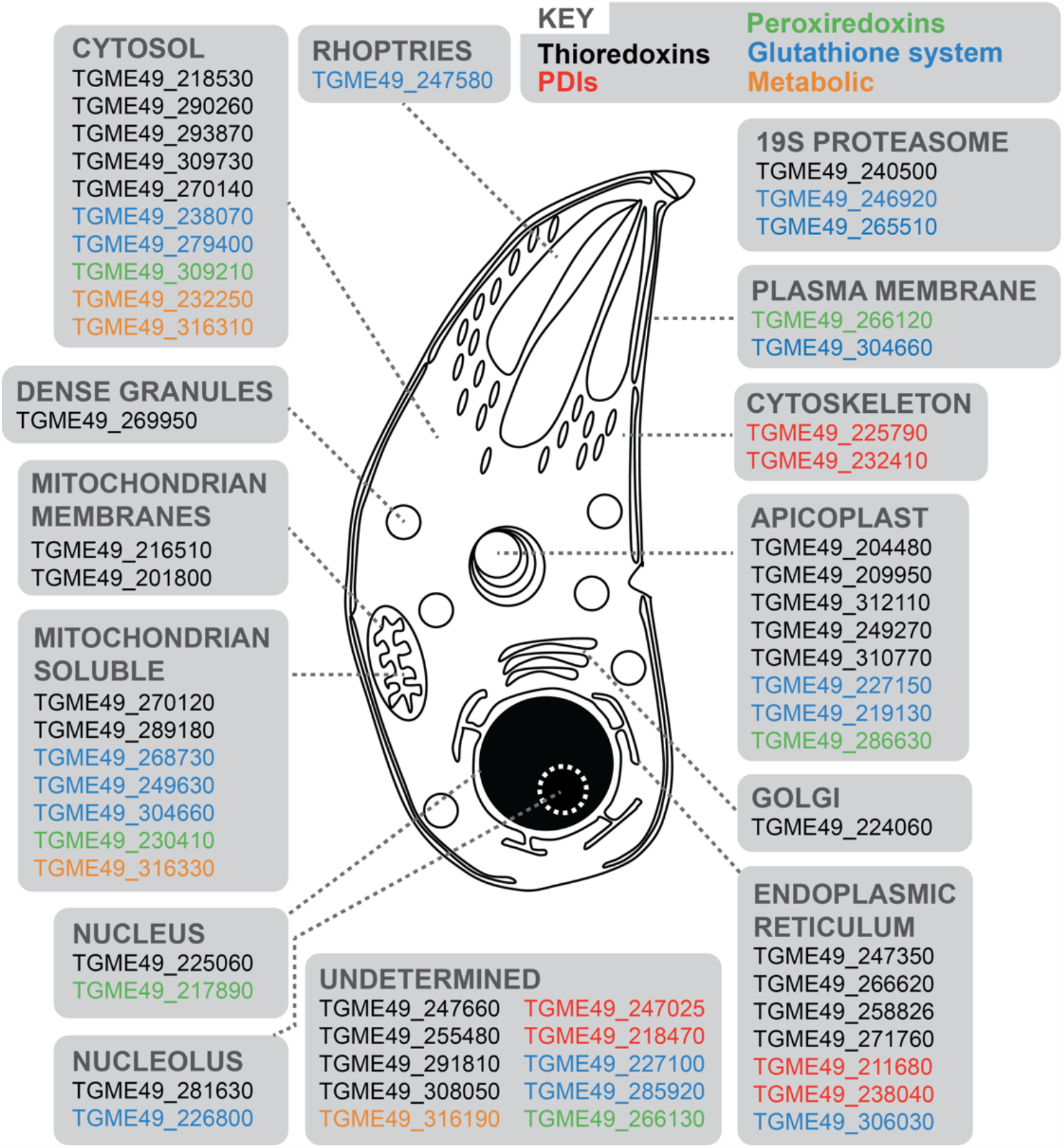
Schematic representation of a *T. gondii* tachyzoite displaying 58 redox-associated genes, and their primary protein location as determined by HyperLOPIT^(38)^. Gene ID accession numbers are provided, and genes categorized into five groups: black (Trxs); red (PDIs); green (Prxs); blue (Grx-GSHs) and orange (metabolic genes). Further details are provided in Supplement table S1.

### Extracellular hydrogen peroxide induces cytosolic Ca^2+^ flux in intracellular tachyzoites

In other eukaryotic systems there are clear associations between calcium and redox signaling. For example, oxidation of cysteines on the ryanodine receptor stimulates calcium release from intracellular compartments^(39, 40)^. To investigate if a similar overlap exists between these two signaling networks in *T. gondii*, we tested the ability of H_2_O_2_ to trigger a Ca^2+^ response in tachyzoites within a host cell. We generated a Type 1 parasite (RH strain) expressing the Ca^2+^ sensor protein jRCamP1b^(41)^ fused to GFP via the T2A peptide for bicistronic protein expression (referred to as RH-GFP-T2A-jRCaMP1b). The ratio of these two distinct fluorescent proteins provided an elegant tool to distinguish changes in fluorescence due to cell movement from the Ca^2+^ signal (Figure 2 – figure supplement 1A and B). Upon exogenous addition of 100 µM H_2_O_2_ we observed a distinct spike in the intracellular Ca^2+^ reporter signal (Figure 2A – videos supplement 1, 2 and 3). To determine whether the oxidative properties of H_2_O_2_ are responsible for triggering parasite Ca^2+^ release, we repeated the experiment in the presence of the antioxidant α-tocopherol (Figure 2B). Pre-treatment of infected host cell monolayers with 50 µM α-tocopherol abolished H_2_O_2_-stimulated Ca^2+^ release in intracellular parasites. The addition of the Ca^2+^ ionophore A23187 caused an intense Ca^2+^ signal spike in the parasite cytosol in both the presence and absence α-tocopherol (Figure 2A and 2B, respectively), suggesting that the presence of the antioxidant does not interfere with Ca^2+^ stores in intracellular parasites. Vehicle solvent for H_2_O_2_ (water) and for A231877 (DMSO) did not trigger Ca^2+^ mobilisation (Figure 2 – figure supplement 1C and D, respectively).

**Figure 2:**
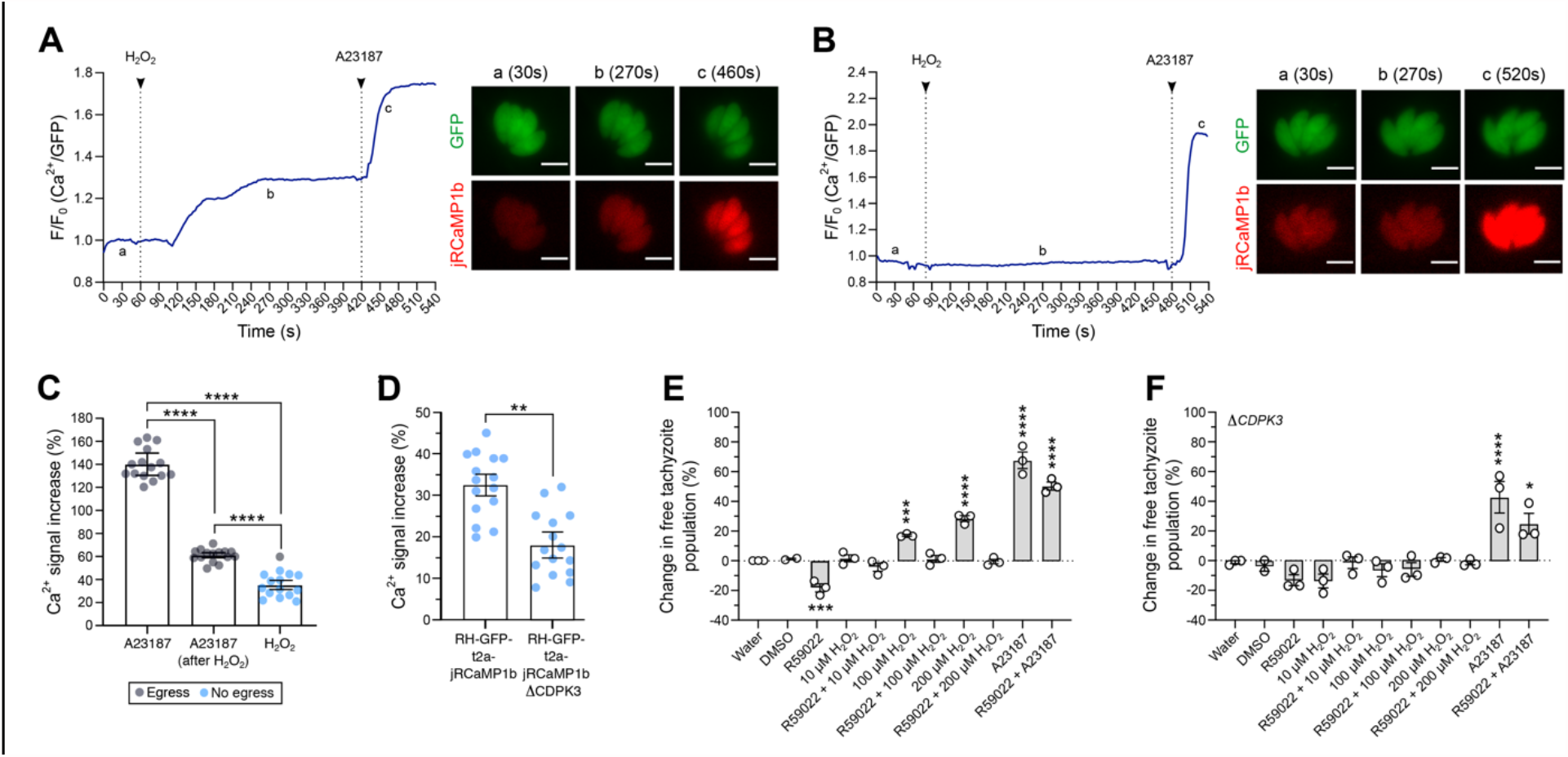
H_2_O_2_ induces Ca^2+^ release in intracellular parasites and triggers CDPK3-dependent egress. Representative trace of 100 µM H_2_O_2_ induction of Ca^2+^ flux in intracellular RH-GFP-T2A-jRCaMP1b parasites. a – c: widefield microscopy images depicting changes in parasite fluorescence signal intensity for both GFP and the Ca^2+^ sensor at baseline (a at 30s), the peak of Ca^2+^ after H_2_O_2_ addition (b at 270s), and peak of Ca^2+^ after 1 µM A23187 addition (c at 460s). Data are representative of 27 infected vacuoles from four independent experiments. (**B**) Representative trace of 100 µM H_2_O_2_ induction of Ca^2+^ flux following pre-treatment of infected host cells with 50 µM α-tocopherol. a: baseline at 30s, b: trace after H_2_O_2_ addition at 270s and <u>c</u>: peak of Ca^2+^ triggered by A23187 at 520s. Data are representative of 14 infected vacuoles from three independent experiments. For (**A**) and (**B**): black arrows indicate the time of compound addition. Scale bar: 5 µM. (**C)** Intensity of parasite Ca^2+^ signal increase in RH-GFP-T2A-jRCaMP1b parasites (expressed as a percentage over baseline) following addition of: 1 µM A23187 alone, 1 µM of A23187 following 100 µM H_2_O_2_ pre-treatment, and 100 µM H_2_O_2_ alone. Gray dots indicate vacuole data points where parasite egress was observed during the measurement period, blue dots represent vacuoles where egress was not observed. (**D)** Intensity of parasite Ca^2+^ signal increase (%) over the baseline upon addition of 100 µM H_2_O_2_ in RH-GFP-T2A-jRCaMP1b *versus* RH-GFP-T2A-jRCaMP1bΔ*CDPK3* parasites. For (**C**) and (**D**): histograms present data mean ±SEM of three independent experiments (five vacuoles measured in each experiment), with individual vacuole data points also shown. (**E**) and (**F**): Egress assay measuring tachyzoite release after compound treatment in RH-GFP-T2A-jRCaMP1b and RH-GFP-T2A-jRCaMP1bΔ*CDPK3*, respectively. Data represent the mean ±SEM of three independent experiments (except for DMSO that has two independent experiments), with six technical replicates for each. All data were normalised to the water control. Significance was calculated using one-way Anova, Bonferroni’s multiple comparisons test. P values: *< 0.05; **< 0.01, ***< 0.001 and ****< 0.0001.

We compared the magnitude of the Ca^2+^ signal increase following H_2_O_2_ treatment with the increase triggered by A23187, a small molecule known to induce parasite egress by Ca^2+^-dependent mechanisms^(42)^ (Figure 2C). H_2_O_2_ induced an overall Ca^2+^ signal increase of 35% ± 1.7. This was lower than the Ca^2+^ signal increase observed when parasites were treated with A23187 alone (140 ± 4.3), or A23187 on cells pre-treated with 100 µM H_2_O_2_ (61% ± 0.9). Within restricted timeframe of this experiment (10 minutes), a single egress event was observed for cells treated with 100 µM H_2_O_2_ (Figure 2C – video supplement 2). Observation of egress induced by H_2_O_2_ was more frequent when host cells were infected with a high multiplicity of infection (MOI = 5) (Figure 2 – video supplement 3). We also measured the magnitude of Ca^2+^ signal increase induced by H_2_O_2_ in Δ*CDPK3* parasites (Figure 2D). The magnitude of the Ca^2+^ signal induced by H_2_O_2_ in RH-GFP-T2A-jRCaMP1bΔ*CDPK3* was lower (18.1% ± 4.3) compared to RH-GFP-T2A-jRCaMP1b (32.5% ± 1.1). This could reflect previously observed differences in the resting Ca^2+^ levels of these lines^(43)^. Together, these data suggest that tachyzoites within the PV can perceive and respond to oxidation events initiated outside the host cell.

### Hydrogen peroxide induces parasite egress in a mechanism dependent on CDPK3

Calcium flux accompanies natural egress^(44)^, and calcium ionophores are well-characterized inducers of egress^(13)^. To better investigate the potential of H_2_O_2_ to induce parasite egress, we analysed populations of infected host cells by flow cytometry. The GFP signal from parasites expressing the Ca^2+^ sensor and particle size were used to distinguish infected host cells from free tachyzoites (Figure 2 – figure supplement 2A). We tested whether treatment of infected host cells with different concentrations of H_2_O_2_ could increase the proportion of free tachyzoites. Incubation of infected host cells with H_2_O_2_ resulted in a dose-dependent increase in the number of free tachyzoites (Figure 2E). We sought to understand how the H_2_O_2_ stimulated egress integrates into our current molecular understanding of this process. The small molecule R59022 is an inhibitor of diacylglycerol kinase^(45)^ that inhibits egress by disrupting the formation of phosphatidic acid^(17)^. For all concentration tested, treatment of cells with H_2_O_2_ resulted in a pharmacological rescue of R59022 egress inhibition. However in the presence of R59022, H_2_O_2_ did not stimulate egress above the baseline value. This could relate to the discrete points of activity for the two targets of R59022: diacylglycerol kinase 1 and 2^(46)^ relative to where the H_2_O_2_ effect feeds into the system. The stimulation of egress by H_2_O_2_ was abolished in Δ*CDPK3* parasites (Figure 2F), demonstrating a dependency upon this kinase similar to other egress agonists such as A231287. The incubation of non-infected HFFs with different concentrations of H_2_O_2_ did not alter the proportion of events in this population (Figure 2 – figure supplement 2B), confirming that the concentrations of H_2_O_2_ used in this protocol did not lyse the host cells.

### Intracellular parasites expressing GRX1-roGFP2 sensor perceive oxidation induced by H_2_O_2_

After confirming that exogenous addition of H_2_O_2_ stimulated Ca^2+^ flux in intracellular *T. gondii* parasites, we investigated the redox state within the parasite cytosol and PV. We reasoned that if calcium release was the direct result of an oxidative signal, the redox status of cellular compartments separating intracellular parasites from the extracellular environment should also be affected. To test this, we generated transgenic parasite strains constitutively expressing the redox sensor protein GRX1-roGFP2, in two different cellular compartments: the parasite cytosol, or targeted to the PV as a consequence of an N-terminal fusion with the GRA8 signal sequence^(47)^. GRX1-roGFP2 is a ratiometric redox reporter that detects changes in both reduced (GSH) and oxidized (GSSG) glutathione^(48)^ (Figure 3A and B). Importantly, the redox relay system underpinning GRX1-roGFP2 is not affected by pH, which is known to confound data interpretation with other redox sensor proteins^(49)^. We used GRX1-roGFP2 parasites to track dynamic changes in GSH/GSSG by fluorescence microscopy, and used the normalized GSH/GSSG signal ratio to measure the intensity of oxidation events (Figure 3C – figure supplement 3A). Intracellular parasites expressing the redox sensor targeted to the PV (RH-GRA8-GRX1-roGFP2) or cytosol (RH-GRX1-roGFP2) detected an oxidation event upon exogenous addition of 100 µM H_2_O_2_ (Figure 3D and E, respectively). The oxidation event within the PV was of greater magnitude compared to that detected within the parasite cytosol, indicating that the strength of the oxidative signal was diminished as it crossed the biological membrane separating these compartments. The water control did not affect the redox signal from GRX1-roGFP2 sensor (Figure 3 – figure supplement 3B).

**Figure 3:**
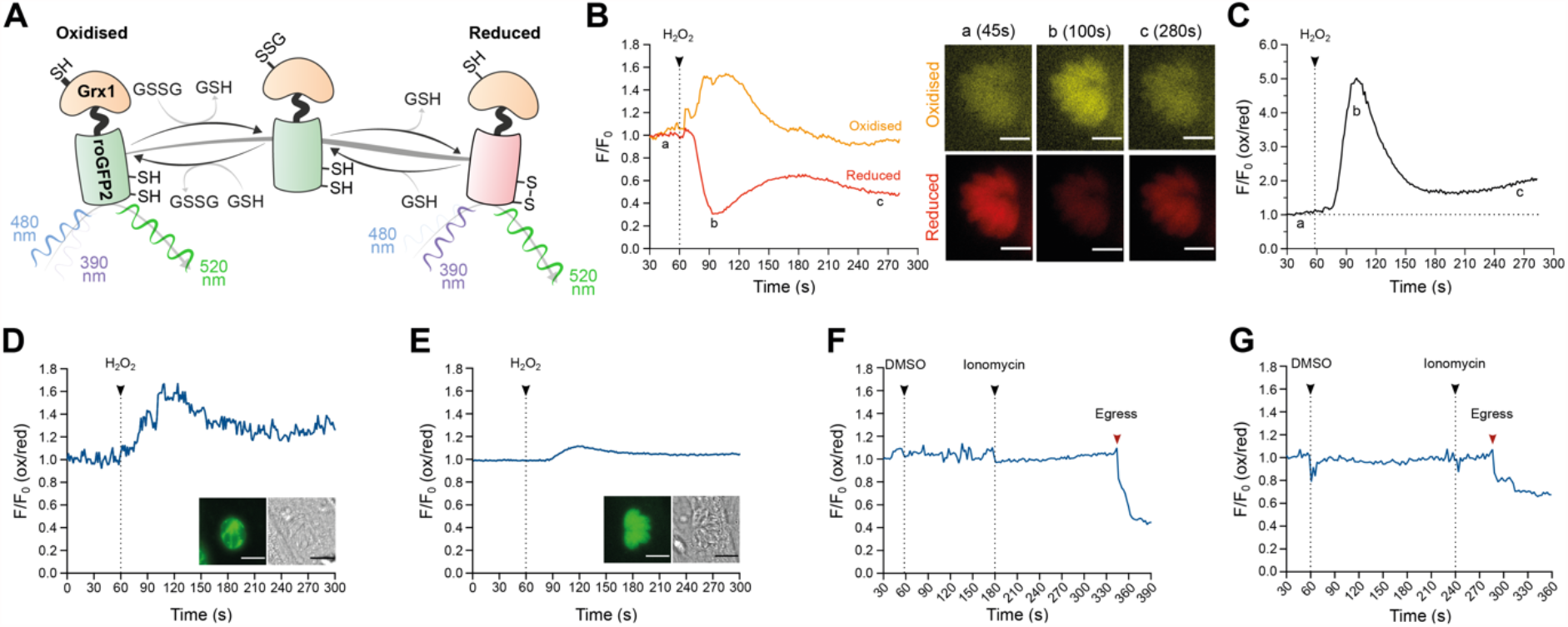
Oxidation induced by exogenous H_2_O_2_ results in a redox change within the PV and parasite cytosol. (**A**) Schematic representation of the redox sensor based on the catalytic domain of glutaredoxin (GRX1) fused with ro-GFP2 depicting how this sensor interacts with GSH/GSSG and the excitation and emission values of the reduced and oxidized states. (**B** – **C**) Proof-of-principle tracking dynamic changes of GSH/GSSG during an oxidation event in RH-GRX1-roGFP2 parasites. Representative signal trace from the GRX1-roGFP2 sensor monitoring both fluorescence channels for reduced (red line) and oxidized (orange line) readouts over time following treatment with 10 mM H_2_O_2_. Microscopy images of an infected vacuole presented in pseudocolor (orange for oxidation, red for reduction) at a: baseline (45s), b: at the peak of the oxidation event (100s) and c: the return to baseline (280s). This is a representative trace from two independent experiments, with eight vacuoles. Scale bar: 5 µm. (**C**) Presents the oxidation/reduction ratio of normalized signal from graph **B** depicting the intensity changes in redox compared to baseline upon addition of 10 mM H_2_O_2_. (**D**) Representative trace from RH-GRA8-GRX1-roGFP2 parasites depicting 100 µM H_2_O_2_ induced redox change within the PV. Microscopy images of an infected vacuole highlighting the presence of the sensor within PV (left image), alongside the brightfield image of the infected host cell. (**E**) A representative trace from RH-GRX1-roGFP2 parasites depicting 100 µM H_2_O_2_ induction of redox change within the parasite cytosol. Microscopy images depicting an infected vacuole highlighting the presence of the sensor on cytosol (left panel image) alongside the brightfield image of the infected host cell. (**D** – **E**): Representative traces from three independent experiment, nine vacuoles for each group. Scale bar: 5 µm. (**F** and **G**): Representative trace of redox fluctuations in intracellular parasite following 1 µM ionomycin treatment for RH-GRA8-GRX1-roGFP2 (**F**), and RH-GRX1-roGFP2 parasites (**G**). (**E** – **F**): red arrows indicate the moment of parasite egress. Representative traces from three independent experiments, nine vacuoles for each group.

Having observed that H_2_O_2_ could stimulate calcium flux, we investigated the reciprocal nature of this relationship by testing the ability of calcium ionophores to trigger an oxidative event. Using GRX1-roGFP2 parasites, we measured the resting redox state prior to ionomycin-induced egress. Ionomycin-induced egress was not accompanied by detectable changes in redox within either the PV or parasite cytosol (Figure 3F and G, respectively). The ionophore A23187 could not be used because of saturating autofluorescence associated with this small molecule in the fluorescence channel used to measure oxidation (Figure 3 – figure supplemental 3C). These data suggested that no significant change in the GSH/GSSG ratio occurs before parasite egress induced by ionomycin.

### GRX1-roGFP2 redox sensor affects parasite fitness during asexual replication

We noted that one unavoidable result of using the GRX1-roGFP2 redox relay sensors would be the associated overexpression of the catalytic domain of glutaredoxin (as a consequence of it being fused to roGFP2). GRX1 is a small redox enzyme that confers protection against oxidative stress^(49, 50)^, and the overexpression of this enzyme would be expected to shift GSH/GSSG ratios in favour of the reduced form (GSH). Correspondingly, we hypothesized that *T. gondii* strains overexpressing GRX1 would have their normal redox status shifted to a more reduced potential. To test whether the overexpression of this redox protein affected parasite asexual growth, we generated a transgenic parasite strain expressing a version of the redox sensor where we had mutated the key catalytic cysteine residue of GRX to render it enzymatically inactive (GRX1_ser_-roGFP2). As a result, GRX1ser-roGFP2 sensor did not respond changes in GSH/GSSG following H_2_O_2_ treatment (Figure 4 - figure supplement 4). We compared the growth of this strain with parasites expressing the catalytically active version of the sensor. As before, both active and inactive versions of the GRX-fusion sensor were targeted to either the parasite cytosol or PV. As a control group, we used a parasite strain expressing redox-insensitive sensor GFP (RH-GFP-Luc)^(51)^. Parasites expressing catalytically active GRX1 in the cytosol presented fewer plaques compared to RH-GFP-Luc (Figure 4A). Despite being able to successfully maintain PV-targeted sensor strains in culture, all parasites lines where the sensor was targeted to the PV did not form clear measurable plaques after six days of growth. Intensely stained plaque-shaped boundaries visible on the HFF monolayer suggested these parasites had successfully grown, but that parasite lytic growth had not outcompeted host cell monolayer recovery sufficiently to produce a clear zone of lysis (Figure 4B). When either the catalytically active or inactive sensors were targeted to the parasite cytosol, parasites formed smaller plaques compared to the RH-GFP-Luc control (Figure 4B and 4C, respectively). These data suggested that the presence roGFP2 alone might be sufficient to affect parasite growth. To directly test the influence of redox environment on the lytic cycle, we grew RH-GFP-Luc parasites with 10 µM N-acetyl cysteine (NAC), a small antioxidant molecule that functions by donating cysteine to increase GSH biosynthesis^(52)^. Addition of NAC decreased the size of plaques generated by RH-GFP-Luc parasites (Figure 4C), suggesting that GSH/GSSG imbalance compromised parasite growth. To better understand the impact of overexpressing the sensor on parasite growth, we counted the number of parasites per vacuole after 20 hours of asexual parasite replication (Figure 4D). All strains expressing the redox sensor had reduced replication compared to RH-GFP-Luc (Figure 4 – figure supplement 5), with catalytic inactivation of the GRX domain providing a partial rescue of the replication defect. Strains exhibiting the slowest replication were those where the sensor was targeted to the PV, supporting the plaque growth data in Figure 4B. Together, these data suggest parasite growth is sensitive to changes in GSH/GSSG ratios.

**Figure 4:**
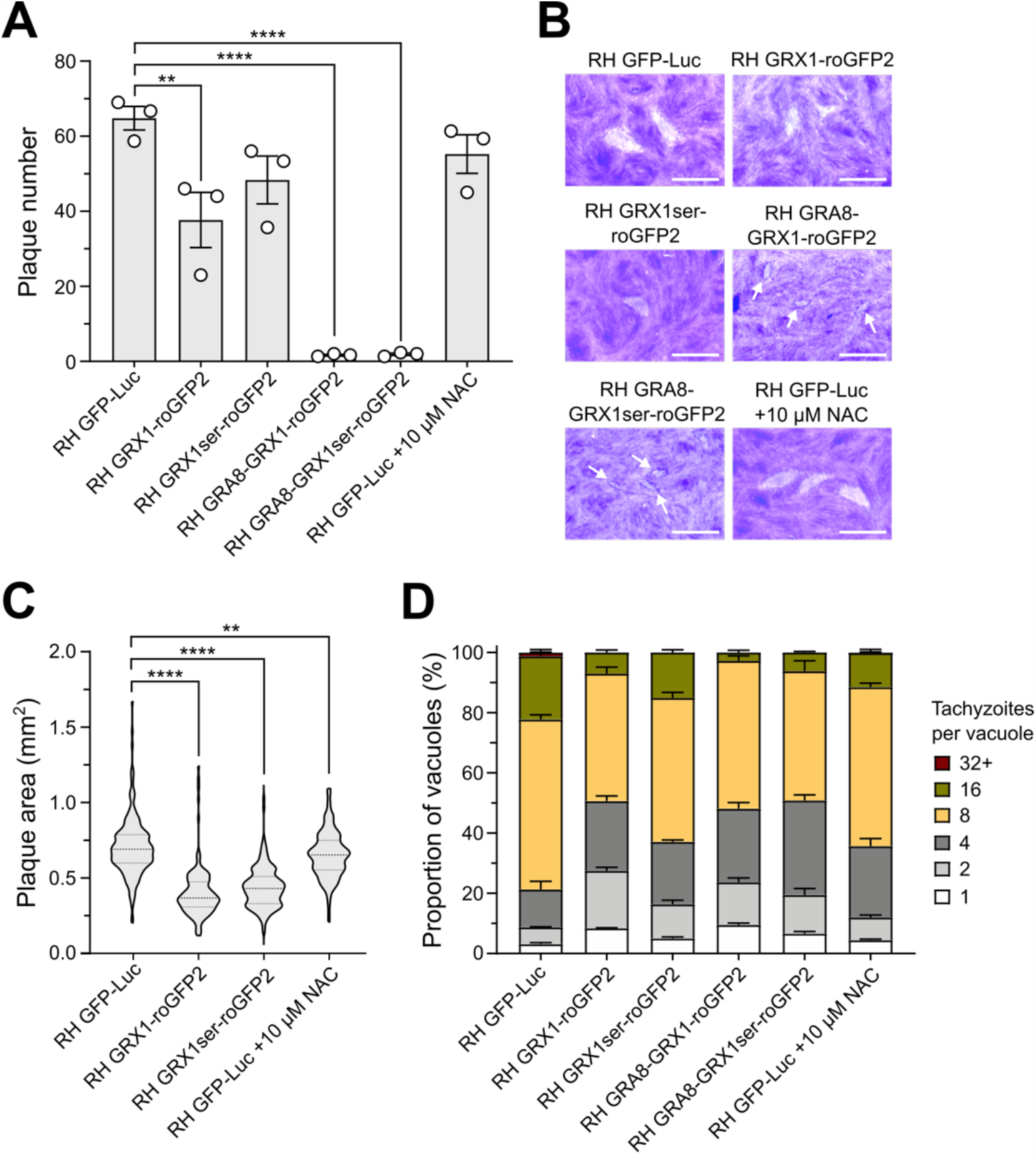
The GRX1-roGFP redox sensor affects *T gondii* asexual replication. (**A**) Histogram presenting plaque count data from a six-day plaque assay using RH-GFP-Luc parasites as a reference control group. (**B**) Representative images of plaques formed. Small plaques are indicated by white arrows. Scale bar: 2 mm. (**C**) Violin plot presenting the distribution of plaque areas (mm^2^) for parasites expressing the redox sensor within the cytosol. (**D**) Histogram presenting the effect of the GRX1-roGFP sensors and NAC on parasite intracellular replication. (**A** – **D**) All obtained from three independent experiments, with three technical replicates. Significance was calculated using one-way Anova, Bonferroni’s multiple comparisons test for (**A**) and (**B**). P values: **< 0.01, **< 0.001 and ****< 0.0001. The significance analyse for (**D**) is provide on figure supplement 5.

## Discussion

Investigations of ROS and *T. gondii* biology initially focused on the host innate immune response. Host macrophages can generate oxidative bursts, creating a toxic microenvironment to fight microbial infection^(53-55)^, with early studies interested in mechanisms used by *T. gondii* to evade oxidative stress^(56-58)^. However, increasing evidence from multiple organisms has demonstrated that H_2_O_2_ has functions as a signaling molecule (reviewed in ^(59)^), suggesting the likelihood of a more complex role for ROS in the pathophysiology of *T. gondii*. To sense, respond, and protect against potential oxidative damage from ROS stimuli, cells employ a network of redox-associated proteins such as Trxs, PDIs, Prxs, Grxs, superoxide dismutase and catalase. It was previously shown that *T. gondii* possesses all these elements^(26)^, and we provide an update on the number, diversity and cellular distribution of these redox-associated proteins (Fig. 1). The function of some of these redox-associated gene products in *T. gondii* parasites has been examined: disruption of catalase decreases parasite virulence in mice and *in vitro* tolerance to H_2_O_2_ ^(26)^; peroxiredoxin-1 interacts with histone lysine methyltransferase to likely regulate gene expression by chromatin rearrangement^(60)^; two thioredoxins have essential roles in apicoplast biogenesis^(61)^, and another thioredoxin with cytosolic localization has been associated with parasite virulence^(62)^. Nevertheless, functional information for most of the redox-associated proteins identified in this work remains elusive.

The ability of low concentrations of H_2_O_2_ to initiate an intracellular Ca^2+^ response in mammalian systems is well documented^(63-65)^ and has helped to solidify H_2_O_2_ as a signaling molecule. Addition of antioxidants blocks intracellular Ca^2+^ release induced by H_2_O_2_ in smooth muscle cells^(65)^ suggesting that the oxidative properties of H_2_O_2_ are required for Ca^2+^ signaling. In human endothelial cells, oxidation induced by non-toxic concentrations of H_2_O_2_ target a Ca^2+^ channel located in acid compartments^(64)^. Distinct from these models, this work is the first to report that an oxidation event induced by H_2_O_2_ mobilizes intracellular Ca^2+^ in *T. gondii* tachyzoites within the infected host cell (Fig. 2A). This is the first time this has been shown for any apicomplexan parasite. Studying the parasite within the host cell provides the best approximation of physiological conditions to observe both Ca^2+^ and redox signaling. Moreover, using parasites expressing a genetically encoded fluorescent Ca^2+^ sensor abrogated the need to use fluorescent Ca^2+^ indicator dyes. Use of these dyes can be damaging to the cells, and typically require the use of other small molecules to avoid dye loss and compartmentalization^(66)^. GFP co-expressed with Ca^2+^ sensor allows Ca^2+^ responses to be distinguished from parasite movement which avoids the need to use inhibitors of parasite motility such as cytochalasin D^(67)^. Finally, the concentration of 100 µM H_2_O_2_ has been showed to be non-toxic for human fibroblast^(68)^. Altogether, our protocol to investigate intracellular Ca^2+^ release following H_2_O_2_ treatment sought to avoid cellular stress that could compromise the true redox/Ca^2+^ dynamic within the parasite. As previously report in muscle cells^(65)^, the presence of antioxidant inhibited the parasite Ca^2+^ response triggered by H_2_O_2_. Using the GRX1-roGFP2 redox sensor, we confirmed that intracellular parasites directly sense an oxidation event following the exogenous addition of 100 µM H_2_O_2_. It is not clear whether the exogenous addition of 100 µM of H_2_O_2_ results in H_2_O_2_ reaching the intracellular parasites as the host cell contains an extensive network of antioxidant proteins that would be expected to scavenge and neutralise H_2_O_2_. This could be directly addressed in future experiments using *T. gondii* strains expressing a sensor to specifically detect H_2_O_2_^(69)^.

Regardless of whether Ca^2+^ is mobilized within the parasite as a direct result of an interaction with H_2_O_2_, or via secondary oxidation signal from the host, Ca^2+^ regulates all aspects of parasite host cell invasion^(9)^ included egress^(16)^. For the short time period used to track parasites by microscopy, the intensity of the Ca^2+^ signal spike induced by H_2_O_2_ was quite modest compared to A23187, and that likely explains why egress events were rare. Longer incubations with H_2_O_2_ induced parasite egress through a mechanism that requires CDPK3 and likely phosphatidic acid. This is the first evidence that suggests that oxidation can trigger *T. gondii* egress. Interestingly, at the other end of the redox spectrum, the reductive molecule DDT can also mobilize Ca^2+^ and induce parasite egress^(31)^. Although the mechanism by which H_2_O_2_ and DDT lead to parasite egress appears to be distinct, these data indicate that redox can influence the parasite’s lytic cycle. We anticipate that parasite biology is tuned to a specific environmental redox potential, and that the perturbation of redox homeostasis with either oxidative or reductive stress elicits a phenotypic response.

The mechanism for how H_2_O_2_ mobilizes Ca^2+^ in animals is better understood but the evolutionary distance between vertebrates and *T. gondii* makes direct comparisons more challenging. *T. gondii* is more closely related to plants, and shares a more similar signaling toolkit^(70)^. It has been recently reported that plants possess a cell surface H_2_O_2_ receptor that once activated, triggers a Ca^2+^ influx into the cells through a Ca^2+^ ion channel^(71)^. *T. gondii* also has a H_2_O_2_-sensitive protein that associates with the CDPK1 to promote microneme secretion^(33)^, an event that requires Ca^2+^ mobilization to allow parasite invasion^(72)^.

Our work suggests a connection between oxidation and parasite Ca^2+^ release but our data did not find evidence that ionophore-induced Ca^2+^ release changes the redox state of either the parasite’s cytosol or PV. The use of the GRX1-roGFP2 sensor to track redox changes based on GSH/GSSG allows a fast and selective assessment of redox fluctuations in real time ^(48)^, with improved dynamics compared to other redox sensors like roGFP1 or roGFP2^(73)^. Considering that Ca^2+^ mobilization is also a fast event, GRX1-roGFP2 is a suitable tool to investigate the relationship between ROS and Ca^2+^. Moreover, the fact that GRX1 does not directly interact with H_2_O_2_ ^(49)^ implies this sensor would not interfere with any eventual interaction between H_2_O_2_ and its potential targets within the parasite or PV. Unexpectedly, expression of GRX1-roGFP2 within *T. gondii* parasites was detrimental to asexual replication. This is surprising as the GRX1-based sensor is generally well tolerated, and presents negligible toxicity in neurons^(49, 74)^. Inactivation of the catalytic domain of GRX1 only partially recovered parasite growth, suggesting that roGFP2 alone is sufficient to influence parasite replication. Cells contain millimolar concentrations of GSH within their cytoplasm and organelles, and under physiological conditions the cell maintains the majority of this redox buffer molecule in a reduced form ([GSH]>[GSSG])^(75)^. Parasites overexpressing GRX1 would be expected to affect the normal GSH/GSSG balance. The addition of NAC, a small molecule that can be used to generate GSH, also slowed parasite growth. This supports the hypothesis that the GSH/GSSG ratio can influence parasite replication. Parasites expressing GRX1ser-roGFP2 are likely have altered redox potential due ability of roGFP2 not fused with an active GRX1 to interact directly with oxidizing molecules like H_2_O_2_^(76)^.

Targeting of the redox sensor to the PV had the greatest effect upon parasite asexual growth. During asexual replication within host cells, *T. gondii* resides within the PV. This compartment separates the parasite from the host cytoplasm, providing a niche for parasite survival and replication^(77)^. Signaling molecules from the host must cross the PV in order to reach the parasite. Should an oxidative signal from the host encounter an unusually reductive environment within the PV due the buffering effect of the redox sensor, the signal could be lost before reaching the parasite. Our work increases our understanding of the complex redox system in *T. gondii*, and provides the first evidence that parasite replication is sensitive to redox imbalance. Redox clearly has an important role in *T. gondii* pathophysiology and we anticipate that the unveiling of this network will guide future covalent drug discovery targeting redox-sensitive chemically reactive cysteines.

## Material and Methods

### Parasite and host cell culture

*T. gondii* tachyzoites from strain RH (Type I) lacking hypoxanthine-guanine phosphoribosyl transferase gene (HXGPRT) were cultivated *in vitro* on monolayers of primary human foreskin fibroblast (HFF, ATTC®) in a humidified incubator at 37°C, 5% CO_2,_ 3% O_2_ atmosphere and maintained in Dulbecco’s Modified Eagle medium (DMEM) supplemented with 10% of foetal bovine serum (FBS) and 2mM L-glutamine, without antibiotics. All culture were tested against *Mycoplasma* infection on a monthly basis.

### Generation of plasmids and transgenic parasites

All primers used in this study are listed in table supplement 2. The GRX1-roGFP2 sensor^(48, 78)^ was amplified from the commercially available vector pEIGW-GRX1-roGFP2 (Addgene plasmid nº64990) using primers 1/2 and insert into digested (HF-EcoRI & PacI) pTUB8 vector containing selectable marker for HXGPRT using Gibson Assembly® Master Mix. The same strategy was used for GRA8-GRX1-roGFP2, using primers 2/3. To generate vectors with the inactivated GRX1 catalytic domain (GRX1_cys23-26_)^(79, 80)^, primers 4/5 were used to replace cysteine with a serine on pTUB8::GRX1-roGFP2 and primers 4/6 on pTUB8::GRA8-GRX1-roGFP2, followed circularization with KLD reaction mix (NEB), resulting the vectors pTUB8::GRX1_(ser23-26)_-roGFP2 and pTUB8::GRA8-GRX1_(ser23-26)_-roGFP2. All vectors were linearized and transfected into RH*Δku80ΔHXGPRT* parasites as previously described^(81)^. Transfected parasites were selected 24 hrs post-transfection by addition of mycophenolic acid (MPA; 25µg/mL) and xanthine (XAN; 50 µg/mL) to culture medium. Strains were cloned by limiting dilution into 96 well plates, and five clones selected. Genomic DNA was extracted from extracellular tachyzoites using Monarch^®^ Genomic DNA Purification Kit (New England BioLabs). Presence of GRX1-based sensors was confirmed using primer pair 7/8.

To generate the calcium sensor construct pUPRT::GFP-T2A-jRCaMP1b, sequence encoding the red fluorescent calcium sensor protein jRCaMP1b^(41)^ was ordered from IDT as a custom synthetic gene, and PCR amplified with appropriate Gibson overhangs using primers 9/10. The 5’UTR of *GRA1* was PCR amplified from pTKO2c^(82)^ using primers 11/12, and the GFP-T2A fusion sequence was amplified from an unpublished in-house plasmid using primers 13/14. All three fragments were subsequently cloned by Gibson assembly into PacI-digested UPRT targeting vector pUPRT-HA^(83)^. The resulting construct was linearised using NaeI and transfected into RH *Δku80Δhxgprt* parasites to generate the GFP-T2A-jRCaMP1b calcium sensor line. Transgenic parasites were subjected to 5’-fluo-2’-deoxyuridine (FUDR) selection (5 µM) 24 hrs after transfection. To generate the GFP-T2A-jRCaMP1b Δ*CDPK3* line, the *HXGPRT* casette (flanked by 5’ and 3’ DHFR UTR sequences) was PCR amplified from *pGRA*-HA_HXGPRT^(84)^ using primers 15/16 (introducing 40bp CDPK3 homology regions to the amplified fragment) and co-transfected into RH *Δku80ΔHXGPRT* with pSag1::Cas9-U6::dbl-sgCDPK3. The pSag1::Cas9-U6::dbl-sgCDPK3 vector was generated by inverse PCR amplification of the pSag1::Cas9-U6^(85)^ vector using primer pairs 17/18 and 17/19 to generate intermediate constructs pSag1::Cas9-U6::sg1CDPK3 (comprising sgRNA1) and pSag1::Cas9-U6::sg2CDPK3 (comprising sgRNA2) respectively. Following circularization of both intermediate constructs using KLD reaction mix, a region comprising sgRNA1 was PCR amplified with primers 20/21 from pSag1::Cas9-U6::sg1CDPK3 and Gibson assembled into Kpn1/XhoI linearised pSag1::Cas9-U6:: sg2CDPK3 to generate the double sgRNA plasmid pSag1::Cas9-U6::dbl-sgCDPK3. Recombinant parasites were selected 24 hrs post transfection as previously described for GRX1-roGFP2. Integration of the HXGPRT cassette at the CDPK3 locus was confirmed using primer pairs 22/23 and 24/25 to confirm 5’ and 3’ integration respectively. Absence of the endogenous CDPK3 locus was confirmed using primers 26/27. RH-GFP-Luc parasites (expressing GFP mutant 3 and firefly luciferase IAV^(51)^) were a gift from Dr Moritz Treeck.

### Identification of *T. gondii* genes related to redox

Within the *Toxoplasma* database (ToxoDB^(35)^, release 50 beta 17 Dec 2020), a list of genes related to redox signaling in *T. gondii* were obtained using the keywords “thioredoxin”, “glutathione”, “glutaredoxin”, “peroxiredoxin” and “protein disulfide” on gene text search option. The function domains of the canonical antioxidant groups Trxs, Grx-GSH, Prxs and PDIs were confirmed for each gene using Basic Local Alignment Search Tool (BLAST) at National Center for Biotechnology information (NCBI) database (www.ncbi.nlm.nih.gov/). The spatial protein localization related to each redox gene were extracted from the localisation of organelle proteins by isotope tagging (hyperLOPIT)^(38, 86)^ dataset available through ToxoDB.

### Plaque and replication assays

For plaque assays, tachyzoites were harvested from infected HFF by syringe passage followed by filtration (5µm). ∼100 tachyzoites were added per well of a 6-well plate prepared with confluent HFF monolayers, and allowed to grow undisturbed for 6 days. Plates were washed with phosphate-buffer saline (PBS) and fixed with cold methanol, stained with crystal violet and scanned. Plaque counts and area measurements were performed using FIJI software by drawing region of interesting (ROI). For replication assays, the freshly lysed tachyzoites were added to HFF monolayers grown on a µ-slide 8 well glass bottom chamber (Ibidi®) with multiplicity of infection (MOI) 1. To synchronize the infection, parasites were allowed to settle onto chilled host cells for 20 minutes, and then allowed to invade for 2 hours at 37°C. Cells were washed with PBS to remove extracellular tachyzoites. Cells were incubated for a further 18 hours, and subsequently fixed with 3% paraformaldehyde at room temperature (RT) for 20 min, and blocked with PBS supplemented with 2% FBS. Parasites per vacuole were counted by excitation with 470nm laser (4% intensity) widefield Nikon Eclipse Ti-E inverted microscope equipped with an ORCA-Flash4.0 camera (Hamamatsu, Japan) and NIS-Elements Viewer software (Nikon), 60x-oil objective. All parasite strains were genetically encoding for GFP or roGFP2. All strains were tested tree independent times, each with three technical replicates.

### Egress assay

Freshly lysed tachyzoites from RH-GFP-t2a-jRCaMP1b or RH-GFP-T2A-jRCaMP1bΔ*CDPK3* parasites were harvested and inoculated (MOI:1) onto confluent HFF cells grown on 24 well plates and grown for 18 hours. Cells were washed with PBS and the growth medium replaced with phenol red free DMEM without FBS and incubate for a further three hours. Medium was removed and cells were incubated for 30 minutes with PBS plus drugs (A23187, R59022, H_2_O_2_) or vehicle controls (water, DMSO). At the end of incubation, the supernatant of the wells was carefully aspirated and cells were detached by 10 min incubation with 200 µL of RT Accutase™. Accutase-release cell suspensions were fixed with 200 µL of 8% paraformaldehyde (20 minutes, RT). Fixed cells were transferred into falcon 5 mL tube with 35 µm nylon mesh cap. Free fluorescent parasite and infected HFF population were analysed and quantified in a BD LRSFortessa™. A total of 5000 events were collected for each tube. Blue laser (488nm) with 530/30 nm filter was used to detect the GFP signal. An uninfected HFF control was used to assess the effect of each drug treatment on host cell gating. Each drug condition was tested two independent times, each with six technical replicates.

### Fluorescence microscopy

For cytosolic Ca^2+^ fluorometric measurements, RH-GFP-T2A-jRCaMP1b or RH-GFP-T2A-jRCaMP1bΔ*CDPK3* parasites were added (MOI:1) to confluent HFF grown on 8 well glass bottom chamber and allowed to grow for 18 hours. Wells were washed with PBS and media replaced by phenol red-free DMEM medium without FBS, and incubate for a further two hours. Images of live infected HFFs were captured at 37°C, using a 60x-oil objective in the same widefield microscope previously describe on replication assay. GFP (470 nm excitation/ 520 nm emission) and jRCaMP1b (555 nm excitation/ 605 nm emission) with 100 milliseconds acquisition rate signal were collected every second for up to a maximum 10 minutes. Drugs were applied on the wells after one minute of acquisition by pipetting. Image analyses were performed using FIJI software. Raw fluorescence readout (F) for each vacuole on each channel was normalised against the average of the baseline signal before adding the drug (F_0_) using the ratio F/F_0_ bringing the resting baseline value to one. To distinguish intracellular Ca^2+^ oscillation signal from vacuole movement, the F_(jRCaMP1b)_/F_0(jRCaMP1b)_ values were normalised against the GFP signal F_(GFP)_/F_0(GFP)_. Ca^2+^ response during a specific time is given by the formula: (F_(jRCaMP1b)_/F_0(jRCaMP1b)_) / (F_(GFP)_/F_0(GFP)_).

For redox measurements, parasites expressing ratiometric GRX1 based sensors were treated as described for RH-GFP-T2A-jRCaMP1b parasites. Oxidized GRX1-roGFP2 (395 nm excitation/ 520 nm emission) and reduced GRX1-roGFP2 (470 nm excitation/ 520 nm emission) with 200 milliseconds acquisition rate signal were collected every second for up to a maximum 10 minutes. Data analyses is similar to Ca^2+^, the redox reading was obtained by formula: (F_(oxidised)_/F_0(oxidised)_) / (F_(reduced)_/F_0(reduced)_).

### Chemical and Reagents

Hydrogen peroxide 30% (w/w) solution with stabilizer; diacylglycerol kinase inhibitor (R59022); A23187, Ionomycin, anhydrous methanol, and crystal violet solution were obtained from Sigma-Aldrich Company Ltd. α-tocopherol phosphate disodium salt from Merk. Dimethylsulfoxide (DMSO) anhydrous and paraformaldehyde 16% solution from Life Technologies. Accutase cell detachment solution from Fisher Scientific Ltd.

### Statistical Analysis

The data are represented as the mean ± SEM and analysed using two-tailed pared Student *t* test between two groups and one-way or 2way analyses of variance (ANOVA) with Bonferroni’s multiple comparisons test for comparing means between ≥3. All data were analysed using GraphPad Prism 9 software (California, USA). The data were considered statistically significant when P values <0.05.

## Funding

This work was supported by grant 202553/Z/16/Z from the Wellcome Trust & Royal Society (to MAC), and BB/M011178/1 from the BBSRC (to HJB and MAC).

## Acknowledgements

We sincerely thank Prof Jake Baum and Dr George Ashdown for the access and assistance to widefield microscopy, Dr Moritz Treeck for the RH-GFP-Luc line, Dr Lilach Sheiner for the roGFP construct, and Dr Gautam Dey for useful discussions. We would also like to thank the Imperial College Flow Cytometry Facility team in South Kensington for technical support. We would like to acknowledge the artistic contribution of Mai Ito who kindly provided the graphical depiction of a *T. gondii* tachyzoite used in Figure 1.

## Competing interests

The authors have declared there are no conflict or competing interest.

## Supplemental Material

**Table supplement 1:**
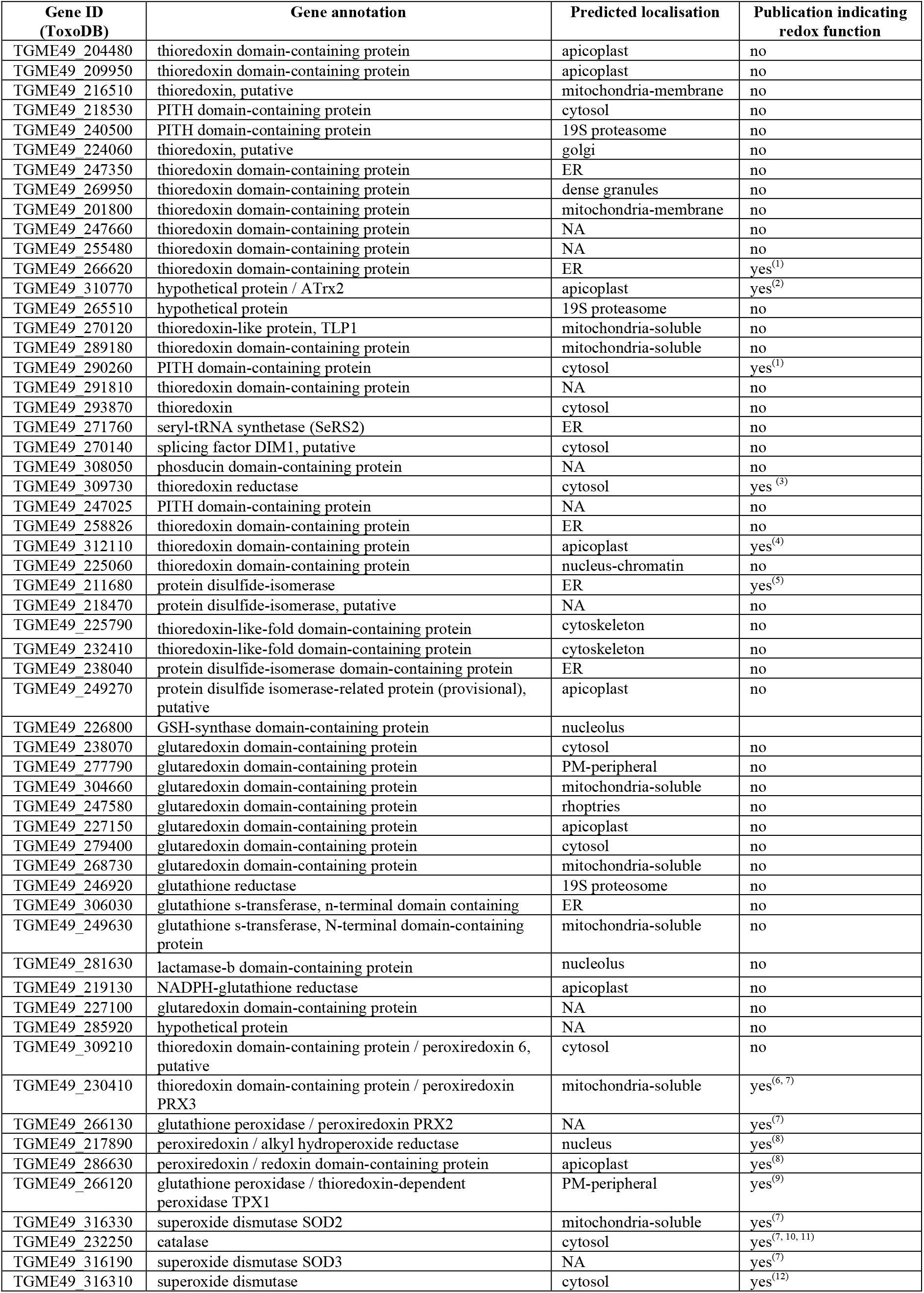
List of *Toxoplasma gondii* genes related to redox sensing systems and their primary location based on hyperLOPIT datasets. Abbreviations: PITH: proteasome-interacting thioredoxin. ER: endoplasmic reticulum. PM: plasma membrane. GSH: glutathione. NA: not applicable (location not predicted).

**Supplemental video 1:** Intracellular *T. gondii* tachyzoites mobilize Ca^2+^ upon addition of H_2_O_2_, followed by egress induced by A23187. The left panel displays the GFP channel, middle panel the Ca^2+^ sensor jCaMP1b channel, and the right panel the channel merge. Time (minutes: seconds) is displayed on the left top region of each channel. 100 µM of H_2_O_2_ was added to cells at 1:06, and 1 µM A23187 was added at 6:51. Video frame rate: 17 frames per second. This video is representative of 27 infected vacuoles from four independent experiments. MOI: 1.

**Supplemental video 2:** Hydrogen peroxide inducing egress of *T. gondii* tachyzoites at low MOI. The left panel displays the GFP channel, the middle panel the Ca^2+^ sensor jCaMP1b channel, and the right panel the merge. Time (minutes: seconds) is displayed on the left top region of each channel. 100 µM of H_2_O_2_ was added to cells at 00:45s. Video frame rate: 10 frames per second. This video presents an egress event captured using low MOI (MOI=1).

**Supplemental video 3:** Hydrogen peroxide inducing egress of *T. gondii* tachyzoites at high MOI. The left panel displays the GFP channel, middle panel the Ca^2+^ sensor jCaMP1b channel, and the right panel the merge. Time (minutes: seconds) is displayed on the left top region of each channel. 100 µM of H_2_O_2_ was added to cells at 00:30s. Video frame rate: 15 frames per second. This video is representative of 34 infected vacuoles from three independent experiments (MOI=5).

**Figure supplement 1:**
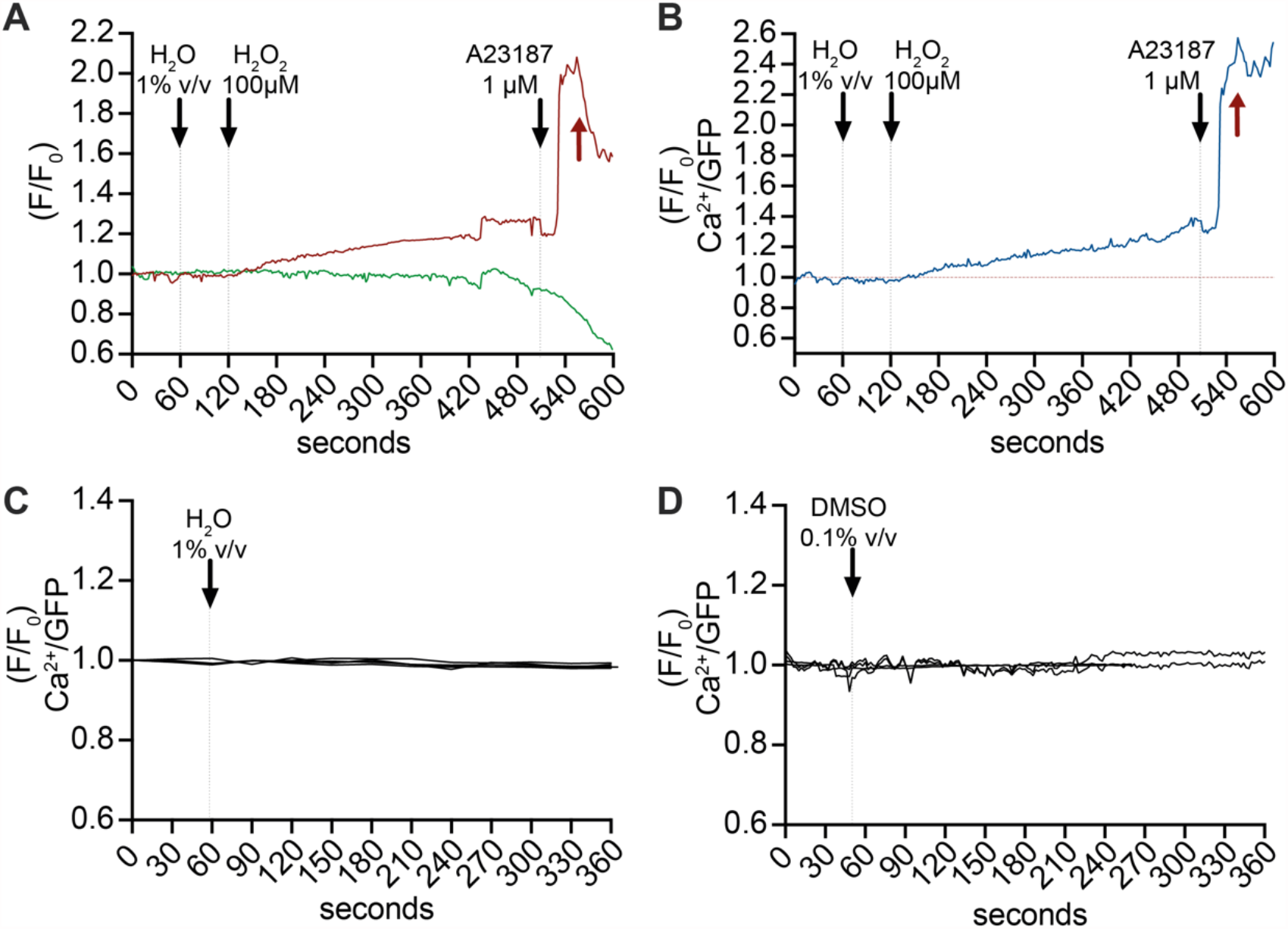
Fluorescence tracking of Ca^2+^ signal and GFP movement on RH-GFP-t2a-jRCaMP1b parasites within host cell. (**A – B**) Representative trace of parasite vacuoles following treatment with H_2_O_2_ and ionophore A23187. (**A**) The graph presents the independent GFP trace (green) and Ca^2+^ signal from jRCaMP1b sensor (red). Note the parasite movement at 420s. (**B**) Graph of the Ca^2+^ signal is normalized to GFP to minimize artefact on the Ca^2+^ measurements due to parasite movement. Red arrow indicates the moment of parasite egress. (**C**) Water, the vehicle solvent for H_2_O_2_, does not mobilize Ca^2+^. The graph displays the trace of three independent vacuoles from the same field of view. Data are representative of 15 infected vacuoles from four independent experiments. (**D**) DMSO, the vehicle solvent for ionophore A23187, does not mobilize Ca^2+^. The graph displays the trace of four independent infected vacuoles from the same field of view. Data are representative of 12 rosettes from three independent experiments. (**A** – **D**): black arrows indicate the time of drug / solvent addition.

**Figure supplement 2:**
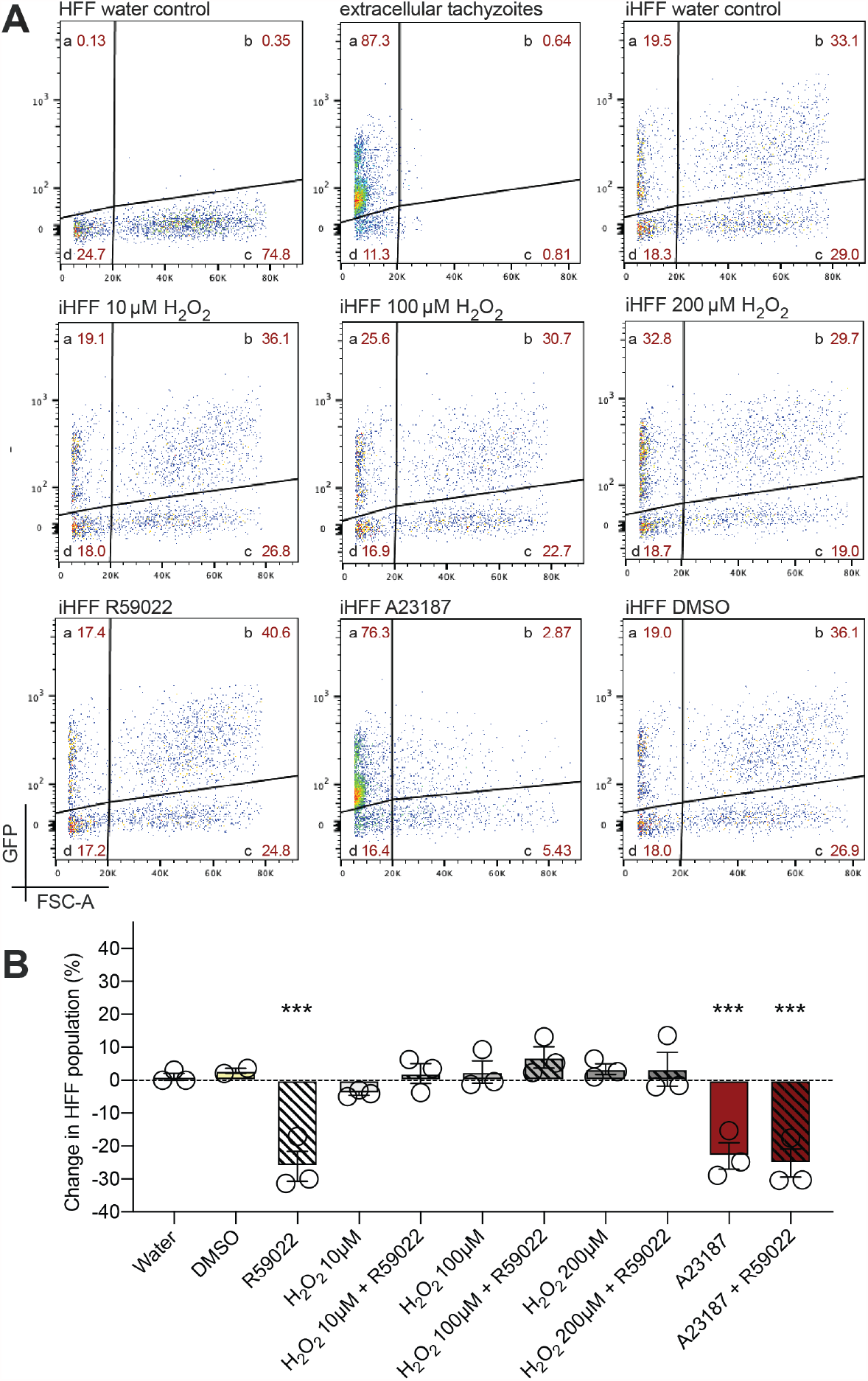
Quantification of RH-GFP-T2A-jCamP1b parasite egress by flow cytometry. (**A**) Representative gating using GFP (Blue laser, 488nm, filter 530/30) against Forward scatter (FSC-A). HFF water control: uninfected human foreskin fibroblast (HFF) are mainly localised in quadrant <u>c</u>. Lysed tachyzoites: free RH-GFP-T2A-jCamP1b parasites are mainly localised in quadrant <u>a</u>. iHFF water control: infected host cells are detected in quadrant <u>b</u>. By using the GFP it is possible to distinguish free fluorescent parasite from iHFF in a population and assess the egress rates throughout different treatments. (**B**) Effect of drug incubation on non-infected HFF. The graph presents a change on event number within the non-fluorescent HFF gate. Data represent the mean ±SEM of three independent experiments (except for DMSO treatment that has two independent experiment), six technical replicates on each. Significance was calculated using one-way Anova, Bonferroni’s multiple comparisons P value ***< 0.001.

**Figure supplement 3:**
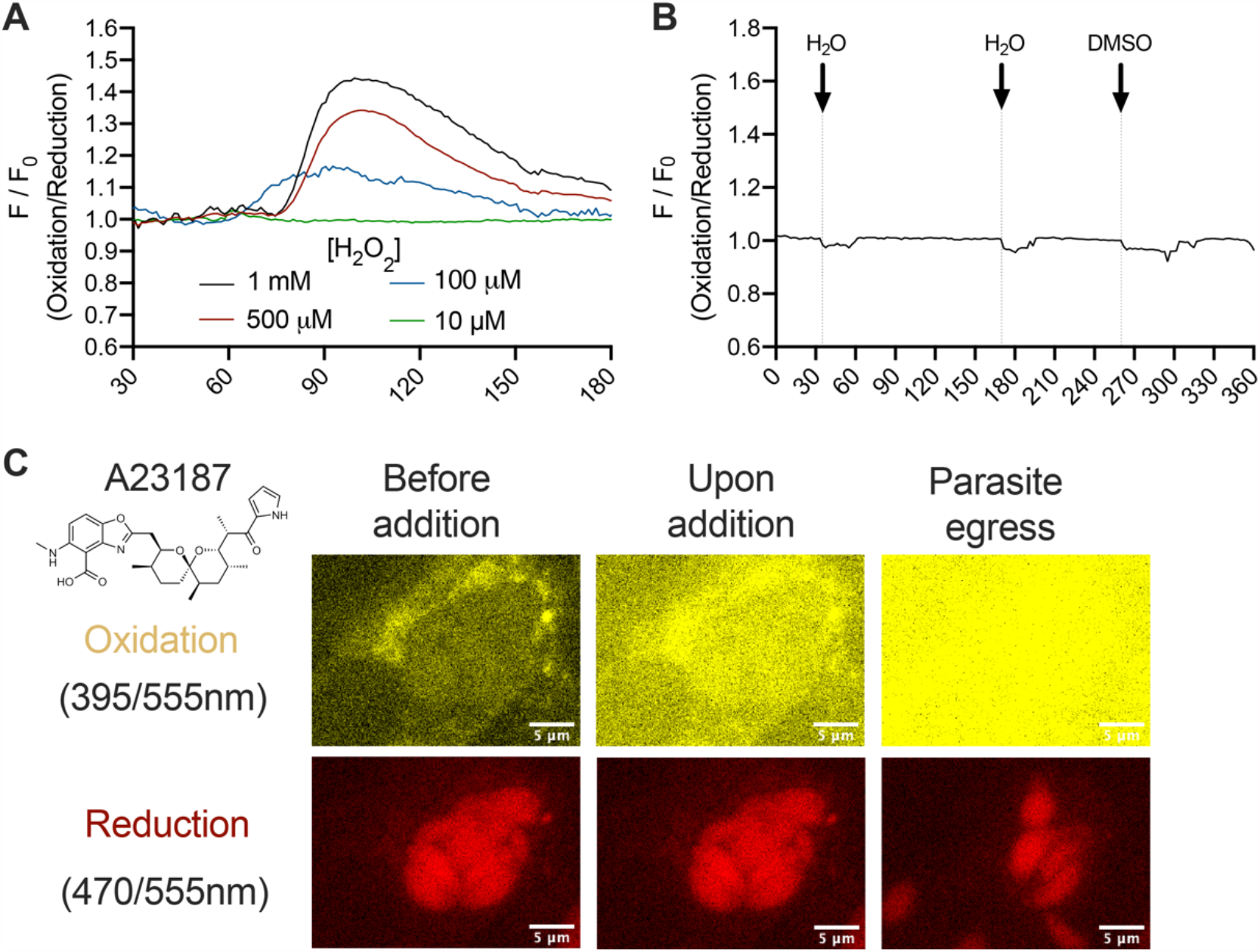
Exploring the redox sensitivity of RH-GRX1-roGFP2 parasites to H_2_O_2_. (**A**) Tracking the GSH/GSSG change upon different concentration of H_2_O_2_. Among the concentration tested, 10 µM was the only one that did not induce a change in redox within the parasite cytosol. Data are representative of five infected vacuoles from each concentration, one independent experiment. (**B**) Water (vehicle control for H_2_O_2_) and DMSO (vehicle control for A23187) do not trigger change in GSH/GSSG. Data are representative of 12 vacuoles from three independent experiments. (**C**) Autofluorescence effect of A23187 drug on oxidation channel makes this ionophore unsuitable for ratiometric analyses of GRX1-roGFP2 sensor. Widefield microscope imaging depicting an egress induce by A23187. The structure of A23187 is shown.

**Supplemental Figure 4:**
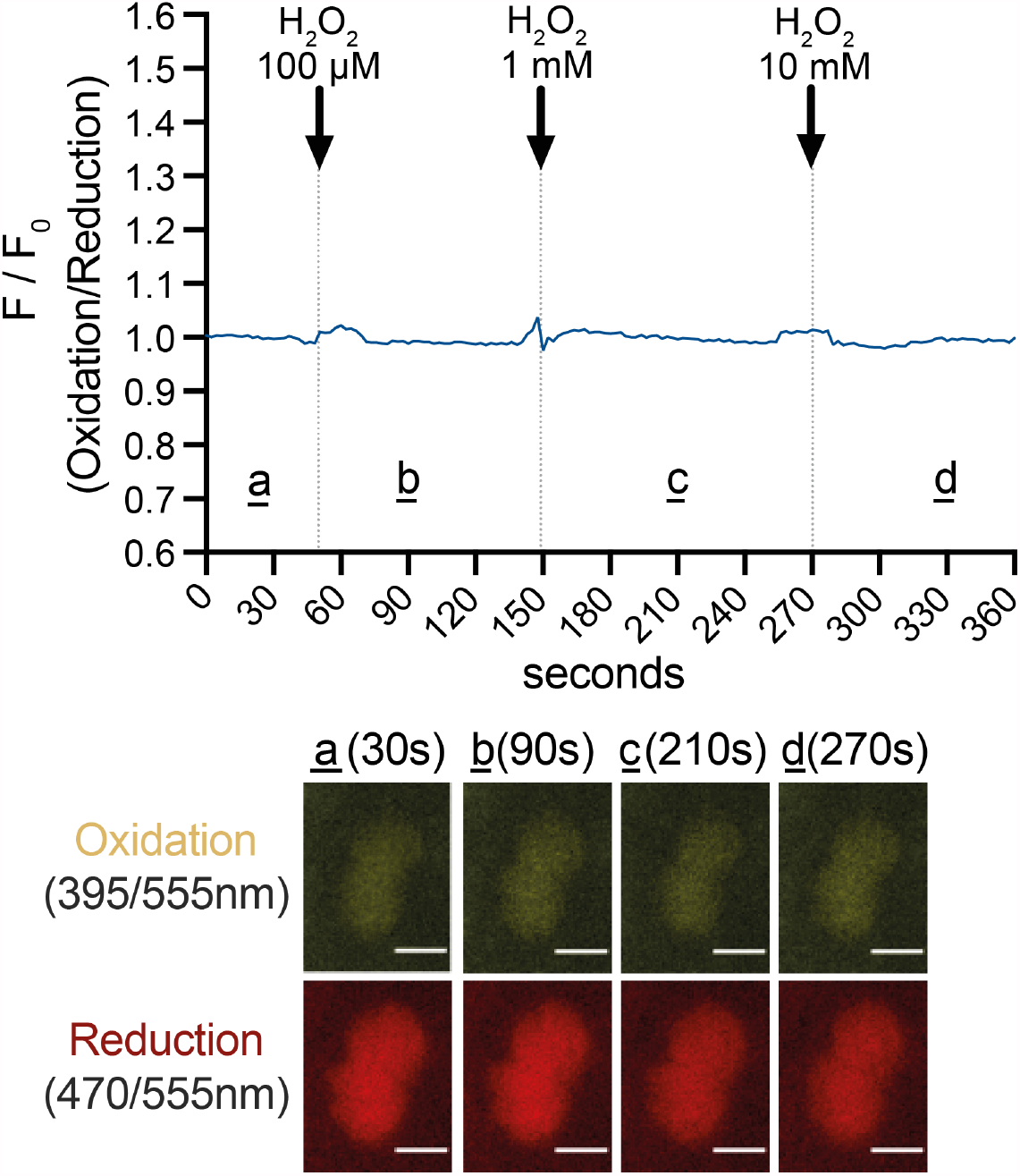
Inactivation of the catalytic domain of glutaredoxin 1 makes the GRX1-roGFP2 redox sensor insensitive to changes in GSH/GSSH. Parasite expressing GRX1ser-roGFP2 do not display fluorescent changes in either channel (reduction or oxidation) upon addition of H_2_O_2_. Data are representative of 12 infected vacuole, three independent experiments. Black arrows indicate the time of H_2_O_2_ addition.

**Figure supplement 5:**
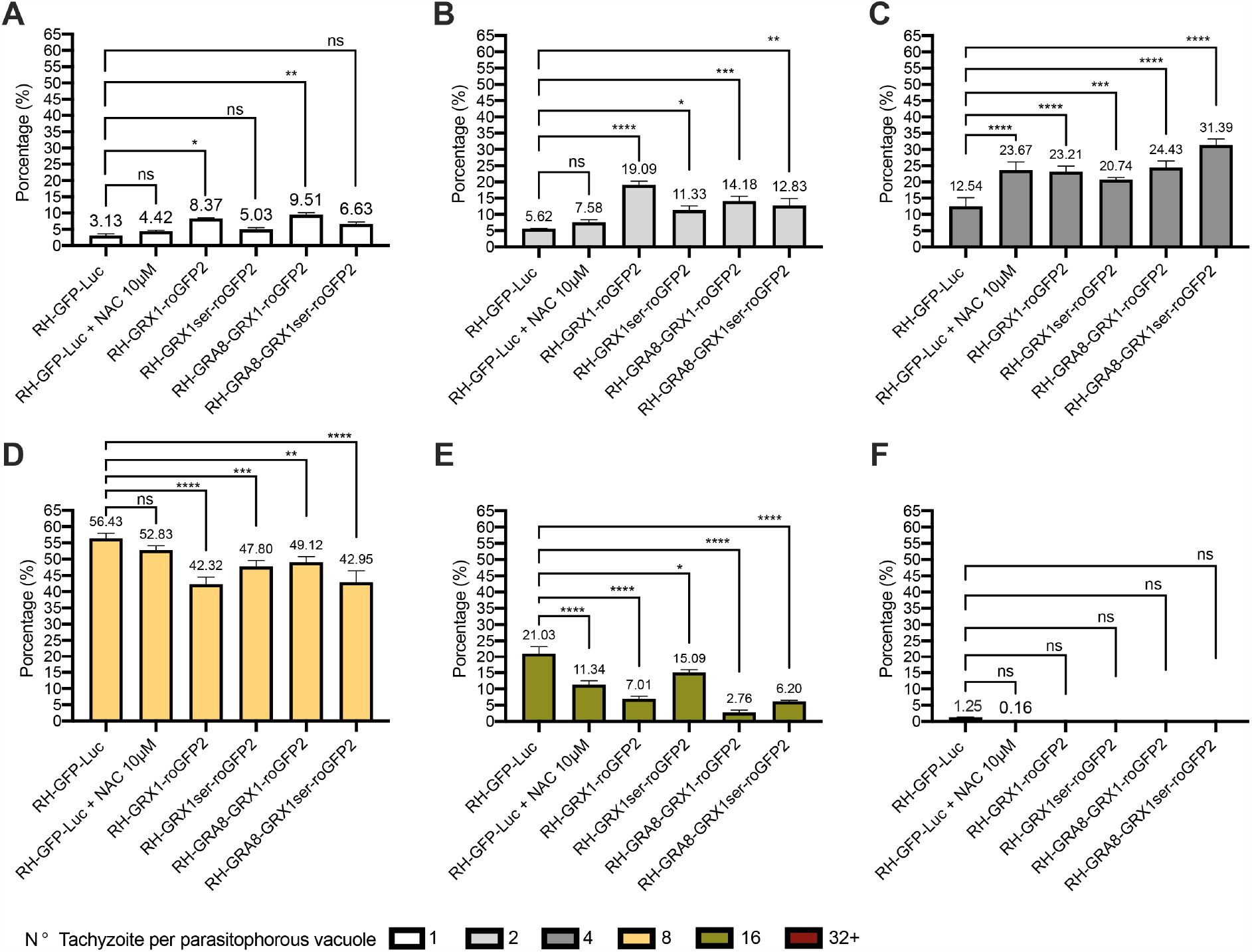
Effect of redox sensors on *T. gondii* parasite growth. Each graph presents parasites/vacuole counts after 20 hours of intracellular growth. The mean values are displayed over each bar for three independent experiments. (**A**) Percentage (%) of vacuoles contain one parasite. (**B**) % of vacuoles with two parasites. (**C**) % of vacuoles with four parasites. (**D**) % of vacuoles with eight parasites. (**E**) % of vacuoles with 16 parasites. (**F**) % of vacuoles with more than 32 parasites. Significance was calculated using two-way Anova, Bonferroni’s multiple comparisons test. P values: *< 0.05; **< 0.01, ***< 0.001 and ****< 0.0001.

**Table supplement 2:**
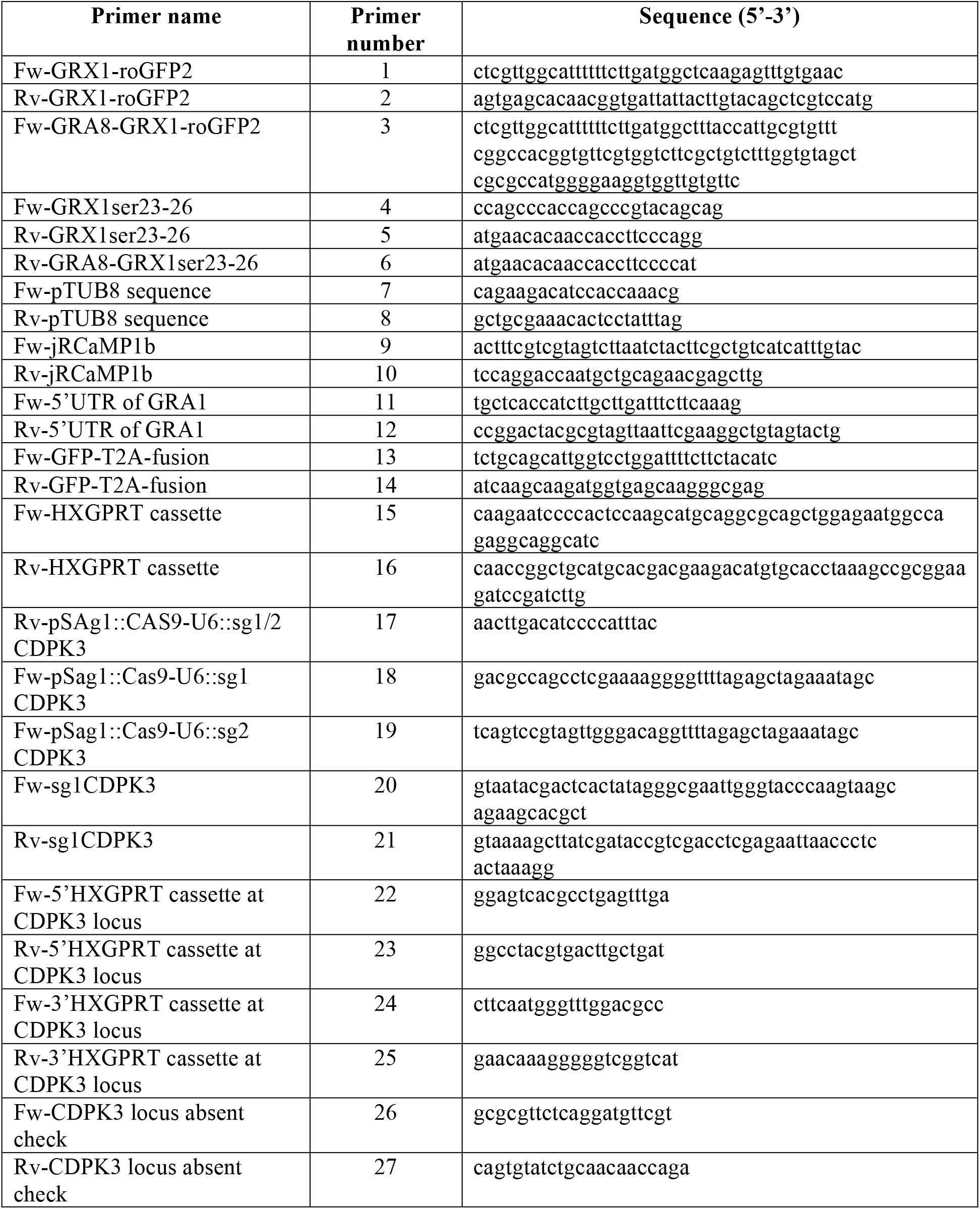
List of primers used to generate plasmid and check sequence / integration in this study.

## Notes

### Competing Interest Statement

The authors have declared no competing interest.

### Summary of Updates

Data descriptions clarified

